# BCCWJ-Brain: A Multi-Modal fMRI, MEG, and EEG Dataset of Naturalistic Japanese Reading

**DOI:** 10.64898/2026.07.05.736621

**Authors:** Yushi Sugimoto, Masayuki Asahara, Hyeonjeong Jeong, Akitake Kanno, Masatoshi Koizumi, Yohei Oseki

## Abstract

We present the BCCWJ-Brain dataset, a multi-modal neuroimaging resource comprising functional magnetic resonance imaging (fMRI), magnetoencephalography (MEG), and electroencephalography (EEG) data recorded from native Japanese speakers reading newspaper articles from the Balanced Corpus of Contemporary Written Japanese (BCCWJ). Neural data were collected from 112 participants (36 fMRI, 35 MEG, and 41 EEG) as they read twenty newspaper articles presented in a Rapid Serial Visual Presentation (RSVP) paradigm. By providing three complementary neuroimaging modalities collected under identical naturalistic reading stimuli, this dataset provides a cognitive benchmark for computational models such as large language models. The dataset is publicly available on the OpenNeuro platform, offering a valuable resource for neuroscience, natural language processing, and related research fields.

## Background & Summary

The past decade has seen rapid growth in the use of neuroimaging data as a benchmark for evaluating computational models of language, particularly large language models, LLMs, (cf. Caucheteux et al., 2023, Gao et al., 2025, Goldstein et al., 2022, Schrimpf et al., 2021, among others). By comparing the internal representations derived from such models with brain activity recorded during naturalistic language comprehension, researchers have probed both the cognitive plausibility of these models and the computational principles underlying language processing in the human brain (cf. Chen and Sivakumar, 2026, Kumar et al., 2024, Lamarre et al., 2022). This comparison between human brain activity and the output from LLMs has been substantially enabled by the public release of large-scale neuroimaging datasets collected under naturalistic stimuli (e.g., Gwilliams et al., 2023, LeBel et al., 2023, Nastase et al., 2021, see also Table 1).

**Table 1:** Recent multimodal neuroimaging datasets for language processing research.

| Dataset name | Language | Modality | Text resources | References |
| --- | --- | --- | --- | --- |
| *fMRI + MEG + EEG* |  |  |  |  |
| BCCWJ-Brain | Japanese | Reading | BCCWJ | This paper |
| *fMRI + MEG* |  |  |  |  |
| SMN4Lang | Mandarin Chinese | Listening | 60 stories (Renmin Daily) | Wang et al. (2022) |
| Mother of Unification Studies | Dutch | Listening & Reading | Normal & scrambled sentences | Schoffelen et al. (2019) |
| *fMRI + EEG* |  |  |  |  |
| LPP Multi-talker Dataset | Mandarin Chinese | Listening | The Little Prince | Wang et al. (2025) |
| Le Petit Prince Hong Kong | Cantonese | Listening | The Little Prince | Momenian et al. (2024) |
| The Alice Dataset | English | Listening | Alice’s Adventures in Wonderland | Bhattasali et al. (2020) |
| *iEEG + fMRI* |  |  |  |  |
| Open multimodal<br>iEEG-fMRI dataset | Dutch | Watching | Short audiovisual film | Berezutskaya et al. (2022) |
| *MEG only* |  |  |  |  |
| MEG-MASC | English | Listening | Manually Annotated Sub-Corpus | Gwilliams et al. (2023) |
| A 10-hour within-participant<br>MEG narrative dataset | English | Listening | Adventures of Sherlock Holmes | Armeni et al. (2022) |
| *fMRI only* |  |  |  |  |
| A natural language fMRI dataset | English | Listening | The Moth | LeBel et al. (2023) |
| The Little Prince Dataset | English/Chinese/French | Listening | The Little Prince | Li et al. (2022) |
| Narratives | English | Listening | Various spoken narratives | Nastase et al. (2021) |
| BBC’s Sherlock Corpus | English | Listening & Speaking | BBC TV series Sherlock | Chen et al. (2017) |

A critical limitation of most existing datasets, however, is that they record only a single neuroimaging modality, which restricts the research questions that can be addressed. Functional MRI (fMRI) measures blood-oxygen-level-dependent (BOLD) signals with high spatial resolution but poor temporal resolution, making it well-suited to identifying where in the brain language processes occur. Electroencephalography (EEG) measures scalp electrical potentials with high temporal resolution but limited spatial precision, and is ideally suited to characterizing the temporal dynamics of language processing, such as the N400 and P600 event-related potential (ERP) components. Magnetoencephalography (MEG) has advantages in both resolutions: its high temporal resolution matches that of EEG, and its relatively high spatial resolution. Because the information that can be captured depends critically on the modality employed, a comprehensive evaluation of a computational model’s cognitive validity requires data from multiple modalities, a standard that single-modality datasets cannot meet.

Several recent multimodal datasets have begun to address this gap, including the Mother of Unification Studies (MoUS) dataset (Schoffelen et al., 2019) for Dutch, the SMN4Lang dataset (Wang et al., 2022) for Mandarin Chinese, the Alice Dataset (Bhattasali et al., 2020) for English, and the Le Petit Prince Hong Kong dataset (Momenian et al., 2024) for Cantonese (Table 1). However, these datasets predominantly target spoken language comprehension, cover a restricted set of languages, and none simultaneously provides fMRI, MEG, and EEG from the same stimuli in a reading paradigm. Moreover, while existing resources span several languages including Chinese and English, Japanese remains notably absent, despite its typologically distinct properties such as its agglutinative morphology and flexible word order, which make it a particularly valuable target for neurolinguistic investigation.

Here we introduce **BCCWJ-Brain**, a new multi-modal neuroimaging dataset designed to address these gaps. The dataset comprises fMRI, MEG, and EEG recordings from 112 native Japanese speakers who read twenty newspaper articles drawn from the BCCWJ (Maekawa et al., 2014), a widely used and linguistically well-annotated corpus. All three modalities were recorded using identical text stimuli, enabling direct cross-modal comparisons.

The dataset is publicly available at OpenNeuro platform, separated in three dataset, BCCWJ-fMRI (ds007752), BCCWJ-MEG (ds007763), and BCCWJ-EEG (ds007753). These datasets include 112 participants’ data in total, in which each participant read twenty newspaper articles segment by segment, spanning 1,642 phrasal units (or *‘bunsetsu’*) and 229 sentences across approximately 30 minutes of stimuli.

## Methods

### Participants

A total of 124 participants read twenty Japanese newspaper articles from *The Balanced Corpus of Contemporary Written Japanese* (BCCWJ) (Maekawa et al., 2014) during neuroimaging acquisition. Following the exclusion of participants with unavailable or unusable data, the final dataset comprises 112 participants.

Separate participant groups were recruited for fMRI, MEG, and EEG data collection. For fMRI, 42 participants were initially recruited; however, six were subsequently excluded; four due to incomplete data acquisition (sub-00, sub-01, sub-02, and sub-28), one due to stimulus mispresentation (sub-35), and one due to excessive head movement (sub-03), which resulted in a final fMRI sample of 36 young adults (15 females; mean age = 21.28, SD = 1.68). For MEG, 41 participants were initially recruited. Six were excluded; four due to a data splitting issue (sub-05, sub-12, sub-15, and sub-26) and two due to BEM file issues (sub-13 and sub-29), resulting in a final MEG sample of 35 young adults (13 females; mean age = 21.71, SD = 2.75). For EEG, all 41 recruited participants were retained in the final sample (21 females; mean age = 20.45, SD = 2.88). Across all three modalities, participants were right-handed and self-identified as native Japanese speakers. All participants provided informed consent prior to the experiment and were compensated for their time. See Tables 2 and 3 for a summary.

**Table 2:** Participants’ information.

| fMRI |  |  | MEG |  |  | EEG |  |  |
| --- | --- | --- | --- | --- | --- | --- | --- | --- |
| participant ID | age | sex | participant ID | age | sex | participant ID | age | sex |
| sub-04 | 21 | M | sub-01 | 24 | M | sub-01 | 35 | F |
| sub-05 | 20 | F | sub-02 | 23 | M | sub-02 | 20 | M |
| sub-06 | 21 | F | sub-03 | 19 | M | sub-03 | 22 | F |
| sub-07 | 20 | M | sub-04 | 31 | M | sub-04 | 22 | M |
| sub-08 | 20 | M | sub-06 | 31 | M | sub-05 | 20 | M |
| sub-09 | 21 | M | sub-07 | 21 | M | sub-06 | 19 | M |
| sub-10 | 21 | M | sub-08 | 19 | F | sub-07 | 20 | M |
| sub-11 | 20 | M | sub-09 | 20 | F | sub-08 | 18 | M |
| sub-12 | 20 | M | sub-10 | 18 | F | sub-09 | 25 | M |
| sub-13 | 23 | M | sub-11 | 21 | F | sub-10 | 19 | M |
| sub-14 | 19 | M | sub-14 | 24 | M | sub-11 | 19 | F |
| sub-15 | 20 | F | sub-16 | 21 | M | sub-12 | 21 | M |
| sub-16 | 22 | F | sub-17 | 21 | F | sub-13 | 19 | M |
| sub-17 | 20 | M | sub-18 | 22 | M | sub-14 | 19 | F |
| sub-18 | 20 | M | sub-19 | 23 | M | sub-15 | 21 | F |
| sub-19 | 20 | M | sub-20 | 22 | F | sub-16 | 19 | M |
| sub-20 | 23 | F | sub-21 | 22 | M | sub-17 | 20 | F |
| sub-21 | 20 | F | sub-22 | 21 | M | sub-18 | 19 | F |
| sub-22 | 24 | M | sub-23 | 20 | M | sub-19 | 19 | F |
| sub-23 | 22 | M | sub-24 | 21 | M | sub-20 | 21 | M |
| sub-24 | 25 | F | sub-25 | 23 | F | sub-21 | 19 | F |
| sub-25 | 19 | F | sub-27 | 22 | M | sub-23 | 20 | M |
| sub-26 | 21 | F | sub-28 | 19 | F | sub-24 | 20 | F |
| sub-27 | 19 | M | sub-30 | 22 | M | sub-25 | 19 | F |
| sub-29 | 20 | F | sub-31 | 20 | M | sub-26 | 19 | F |
| sub-30 | 26 | M | sub-32 | 22 | M | sub-27 | 21 | M |
| sub-31 | 23 | F | sub-33 | 22 | F | sub-28 | 20 | F |
| sub-32 | 21 | F | sub-34 | 20 | F | sub-29 | 19 | F |
| sub-33 | 21 | M | sub-35 | 19 | F | sub-30 | 20 | F |
| sub-34 | 22 | M | sub-36 | 20 | F | sub-31 | 20 | M |
| sub-36 | 20 | M | sub-37 | 22 | M | sub-32 | 27 | F |
| sub-37 | 22 | M | sub-38 | 22 | F | sub-33 | 21 | F |
| sub-38 | 22 | M | sub-39 | 22 | M | sub-34 | 23 | F |
| sub-39 | 20 | F | sub-40 | 19 | F | sub-35 | 20 | M |
| sub-40 | 24 | F | sub-41 | 22 | M | sub-36 | 19 | F |
| sub-41 | 22 | M |  |  |  | sub-38 | 18 | F |
|  |  |  |  |  |  | sub-39 | 21 | M |
|  |  |  |  |  |  | sub-40 | 21 | F |
|  |  |  |  |  |  | sub-41 | 19 | M |
|  |  |  |  |  |  | sub-43 | 18 | M |
|  |  |  |  |  |  | sub-44 | 19 | F |

**Table 3:**
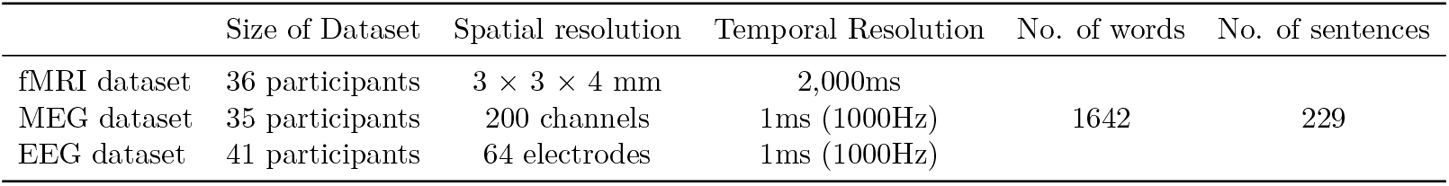
Summary information of the dataset.

### Procedure

#### Stimuli and Procedure

BCCWJ (Maekawa et al., 2014) were used for stimuli. Each newspaper articles were segmented into phrasal units (‘*bunsetsu*’) prescribed by the National Institute for Japanese Language and Linguistics. The stimuli consist of 1642 phrasal units, 229 sentences. The duration of the experiment is around 30 minutes long for reading excluding comprehension questions sessions. Each phrasal unit was presented for 500ms in Rapid Serial Visual Presentation (RSVP) using PsychoPy (Peirce, 2007, 2009), followed by a 500ms blank screen. The presentation order of the articles was randomized for each participants, for each article, one comprehension question was presented and the participants answered the yes-no questions regarding the content of the article by pressing the button on the left index finger if the answer is ‘yes’, or on the left middle finger if the answer is ‘no’.

Before MEG session, we digitized the shape of participant’s faces with a FastSCAN laser scanner (Polhemus), and the five head-position coils. T1-weighted anatomical scans were collected for each participants before or after the MEG session. Likewise, T1-weighted images were collected before or after the fMRI session.

#### Scans

##### fMRI

MRI images were acquired using a Philips Achieva 3.0T MRI scanner. Functional images were collected using an echo-planar imaging pulse sequence (TR = 2,000 ms, echo time = 30 ms, flip angle = 80°, slice thickness = 4 mm, no slice gap, field of view = 192 × 192 mm, matrix = 64 × 64, voxel size = 3 × 3 × 4 mm). T1-weighted anatomical images were acquired from all participants. Scanning parameters included a slice thickness of 1 mm, field of view of 256 × 256 mm, matrix size of 368 × 368, repetition time (TR) of 1,100 ms, and echo time (TE) of 5.1ms. Facial structures were removed from T1-weighted images using PyDeface (Gulban et al., 2022).

##### MEG

Continuous MEG was recorded with a 200-channel whole-head MEG system with axial gradiometers (RICOH Ltd., Tokyo, Japan) in a shielded room at a sampling rate of 1,000 Hz with an online low-pass filter of 200 Hz. T1-weighted images were also obtained before or after the MEG experiments for each participant (the parameters were the same as the one above). T1-weighted images were defaced using PyDeface (Gulban et al., 2022).

##### EEG

EEG data were recorded using a BrainAmp amplifier (Brain Products GmbH, Germany) with a 64-channel electrode cap (including IO, an electrooculogram (EOG) channel for ocular artifact detection). The online reference electrode was placed at FCz, and the ground electrode was placed at AFz. An electrode was placed below the right eye (IO) to monitor ocular artifacts. Electrode impedances were kept below 20 kΩ prior to the recording. Data were recorded at a sampling rate of 1,000Hz.

#### Preprocessing

##### fMRI

MRI files were converted from DICOM to NIfTI formats using dcm2niix.^1^ The preprocessing for functional scans were done using MATLAB (MathWorks, Natick, MA, USA) and Statistical Parametric Mapping (SPM12) software. The preprocessing pipeline included motion realignment, slice timing correction, coregistration to structural images, tissue segmentation, normalization to MNI space, and smoothing with an 8 mm FWHM Gaussian kernel.

##### MEG

The raw MEG data was collected with 1,000Hz online. Continuously Adjusted least Square Method (CALM) filter (Adachi et al., 2001) were applied. Then the continuous MEG data were combined with digitized files and converted into raw.fif files for further analysis. In order to remove ocular artifacts such as eye blink, Independent Component Analysis (ICA) was applied. Then a bandpass filter was applied to 0.1-40 Hz. Data were epoched from -0.1-1,000 ms relative to word onset and baseline correction was applied using the pre-stimulus window of -0.1 to 0ms. For each subject, any channel exceeds a peak-to-peak amplitude threshold of 2e-12 Tesla is rejected automatically. Data were downsampled to 200 Hz for the further analysis to reduce computational load.

##### EEG

The preprocessing procedure for EEG data is similar to MEG preprocessing; ICA was applied for removing artifacts. A 0.1-40Hz bandpass filter was applied. Then, the average of all sensors was used for re-references. Epoching time window was -0.1 to 1,000 ms, with applying baseline correction using the -0.1 to 0.0 ms pre-stimulus window. Epochs that exceed a threshold of 150*µ*V were rejected. Data were downsampled from 1,000 Hz to 200 Hz.

`MNE-python` (Gramfort et al., 2013)^2^ and `Eelbrain` (Brodbeck et al., 2023)^3^ were used for the preprocessing of MEG and EEG data. All preprocessed files are in `derivatives` in the dataset.

### Data Records

The BCCWJ-Brain dataset is publicly available on OpenNeuro. Data are organized according to the BIDS specification (Gorgolewski et al., 2016), including the BIDS extensions for MEG (Niso et al., 2018) and EEG (Pernet et al., 2019).

Each dataset is published separately to make it compatible with the BIDS structure.

- BCCWJ-fMRI: openneuro (https://openneuro.org/datasets/ds007752)
- BCCWJ-MEG: openneuro (https://openneuro.org/datasets/ds007763)
- BCCWJ-EEG: openneuro (https://openneuro.org/datasets/ds007753)

The data structure is represented in Figure 1.

**Figure 1.**
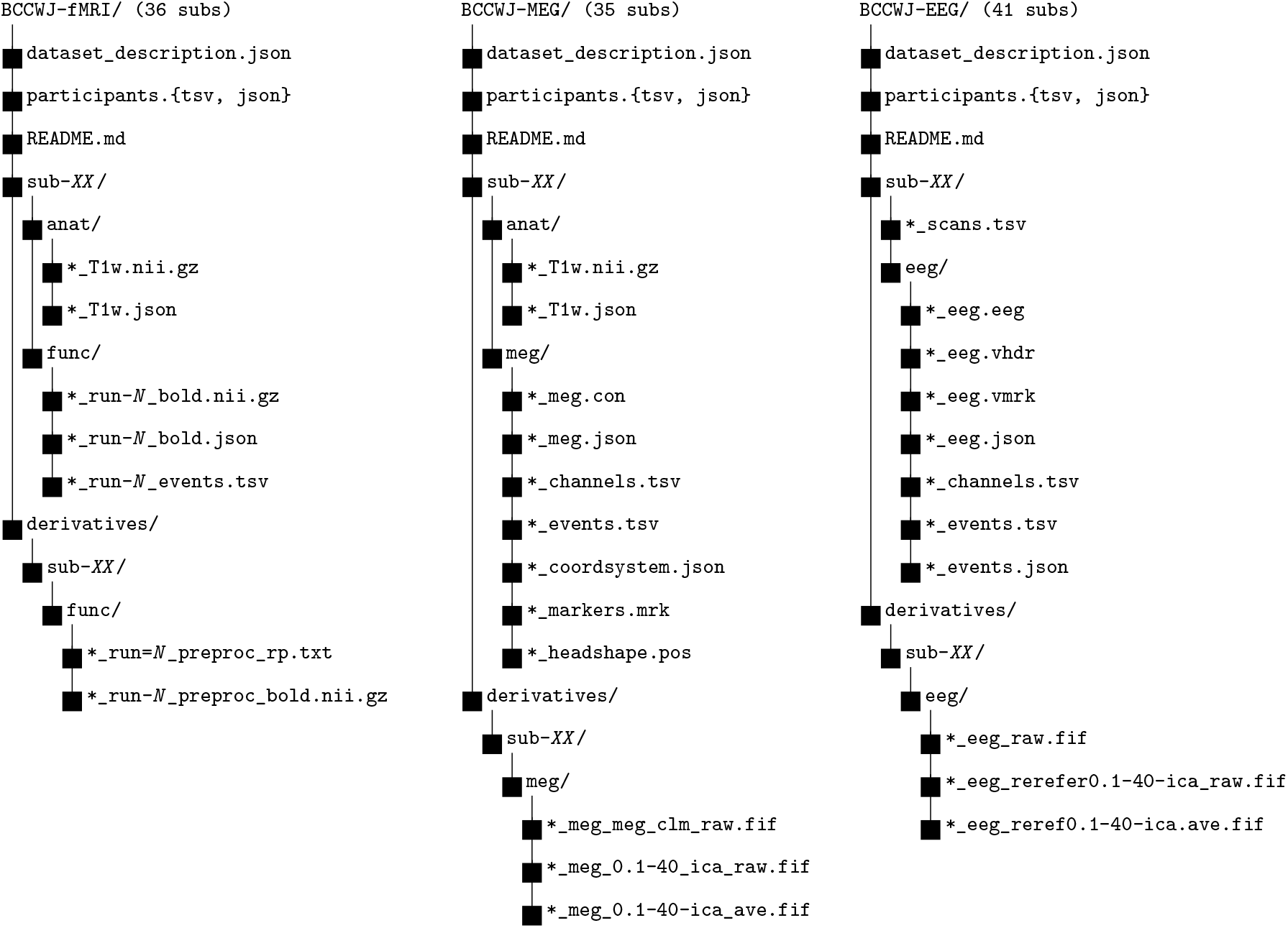
Directory structure of the BCCWJ-Brain datasets (BCCWJ-fMRI, BCCWJ-MEG, and BCCWJ-EEG, which use BIDS structures (v 1.9.0)). *XX* = subject ID; *N* = run number (1–4); * = sub-*XX* _task-BCCWJreading.

**Figure 2.**
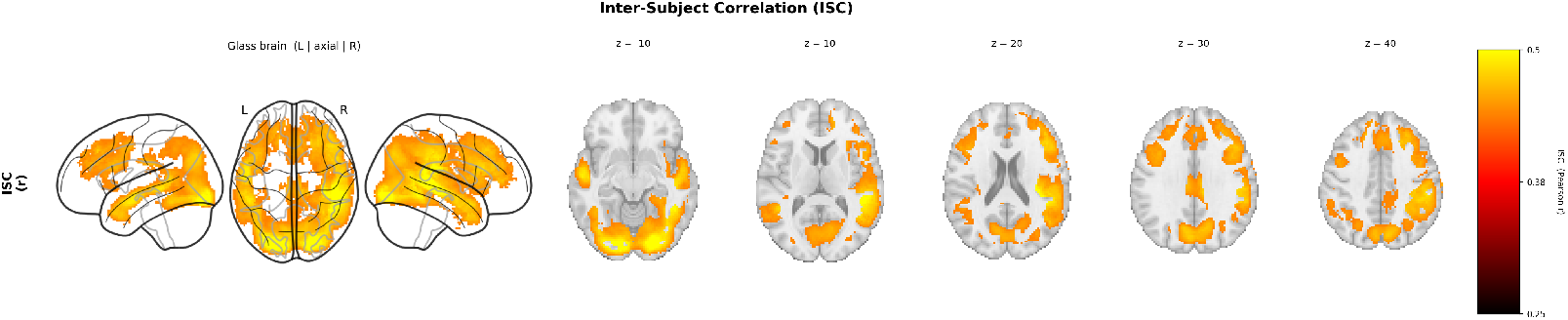
Inter-subject correlation (ISC) results.

**Figure 3.**
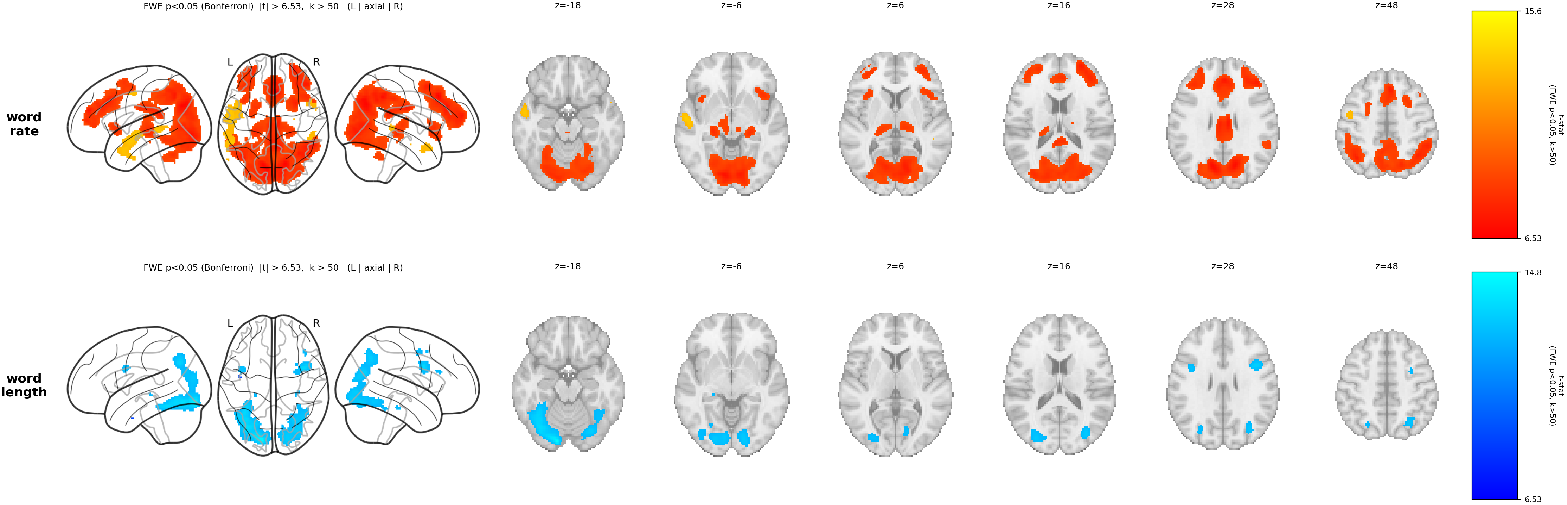
GLM analysis results. Top: word-rate contrast (FWE *p <* 0.05, Bonferroni, *k >* 50 voxels). Bottom: word-length contrast (FWE *p <* 0.05, Bonferroni, *k >* 50 voxels). Color bar indicates *t*-statistic values. L = left hemisphere; R = right hemisphere.

#### Event files

Each functional file is accompanied by a `*_events.tsv` file specifying the onset time, duration, and identity of each presented the phrasal unit (‘*bunsetsu*’). Because the BCCWJ text is not freely available (see Usage Notes), event files only have basic information such as onset time. Researchers with a BCCWJ license can use the provided reconstruction script (see Code Availability) to link event files to the corresponding text.

### Technical Validation

#### Behavioral Results

Participants answered 20 yes-no comprehension questions regarding the content of each newspaper article. For fMRI experiment participants, a mean accuracy for these comprehension questions was 68% (SD = 3.6). For MEG and EEG experiments, the mean accuracies were 81% (SD =7.54) and 85% (SD=6.33) respectively.

#### fMRI

To assess the consistency of neural activity across subjects, we computed inter-subject correlation (ISC) for each voxel using a leave-one-out approach (cf., Nastase et al., 2019). For each participant, we correlated their voxel-wise BOLD time series with the mean time series of all remaining participants. This procedure was applied separately for each of the four experimental sections; the resulting ISC maps were then averaged across sections. At the group level, the median ISC was computed across participants. The results revealed strong correlations across a broad range of regions encompassing the language network, visual cortex, and the visual word form area, suggesting that the observed brain activity across participants reflects language processing engaged through reading.

In addition to the ISC analysis, we validated the fMRI data with the general linear model (GLM) using two regressors of interest; `word rate`, which indicates 1 for the each phrasal unit offset and 0 otherwise. This predictor is expected to show the broad brain activity related to language (Brennan et al., 2012). Another predictor is `word length`, the length of each phrasal unit; since the stimuli were presented on the screen, this predictor activate the brain activity in the visual cortex as well as the visual word form area, which is another expected effect during the scanning. Each predictor was annotated and convolved with canonical hymodynamic response function (implemented in `nilearn`^4^). These predictors were fitted to the fMRI timeseries via GLM for the first level analysis. The first-level effects maps are fed into a one-sample *t*-test with a single intercept regressor. Family-Wise Error (FWE) correction is applied to each group *t*-statistic map.

The results showed that the broad areas of the bilateral temporal lobe (anterior part of the superior temporal gyrus, Table 4) were associated with `word rate`, while the bilateral temporal occipital fusiform cortex (Table 5) were related to `word length`. suggesting that the dataset successfully captures neural responses associated with fundamental aspects of language processing during reading. These results align with existing literature, suggesting that the dataset successfully captures neural responses associated with fundamental aspects of language processing during reading.

**Table 4:** Word rate — FWE (*p <* 0.05, Bonferroni, *k >* 50 voxels, *t >* 6.53).

| Cluster | Voxels | Peak $t$ | $p$ | MNI (x,y,z) | Region |
| --- | --- | --- | --- | --- | --- |
| 1 | 905 | 9.40 | $< .001$ | (-58, -10, -6) | L.Superior Temporal Gyrus, anterior division |
| 2 | 109 | 8.45 | $< .001$ | (48, 16, -24) | R.Precentral Gyrus |
| 3 | 72 | 7.98 | $< .001$ | (50, -34, 2) | R.Superior Temporal Gyrus, anterior division |

**Table 5:** Word length — FWE (*p <* 0.05, Bonferroni, *k >* 50 voxels, *t >* 6.53).

| Cluster | Voxels | Peak $t$ | $p$ | MNI (x,y,z) | Region |
| --- | --- | --- | --- | --- | --- |
| 1 | 3075 | 14.82 | < .001 | (-14, -88, -16) | L.Temporal Occipital Fusiform Cortex |
| 2 | 295 | 10.62 | < .001 | (40, 6, 32) | R.Inferior Frontal Gyrus, pars opercularis |
| 3 | 1186 | 9.46 | < .001 | (32, -76, -16) | R.Temporal Occipital Fusiform Cortex |
| 4 | 832 | 9.30 | < .001 | (34, -70, 22) | R.Angular Gyrus |
| 5 | 68 | 7.94 | < .001 | (-40, 4, 28) | L.Inferior Frontal Gyrus, pars opercularis |

#### MEG

Since there is no standard methods of validating MEG data, we simply show the grand average of the MEG sensor signals, epoching whenever the phrasal unit is presented, averaging the epoching across subjects. See Preprocessing section for the details.

In Figure 4, the sensor level grand average showed the first peak at around 170 to 200ms, which is a typical neural response observed in MEG studies on visual word recognition (Pylkkänen and Marantz, 2003).

**Figure 4.**
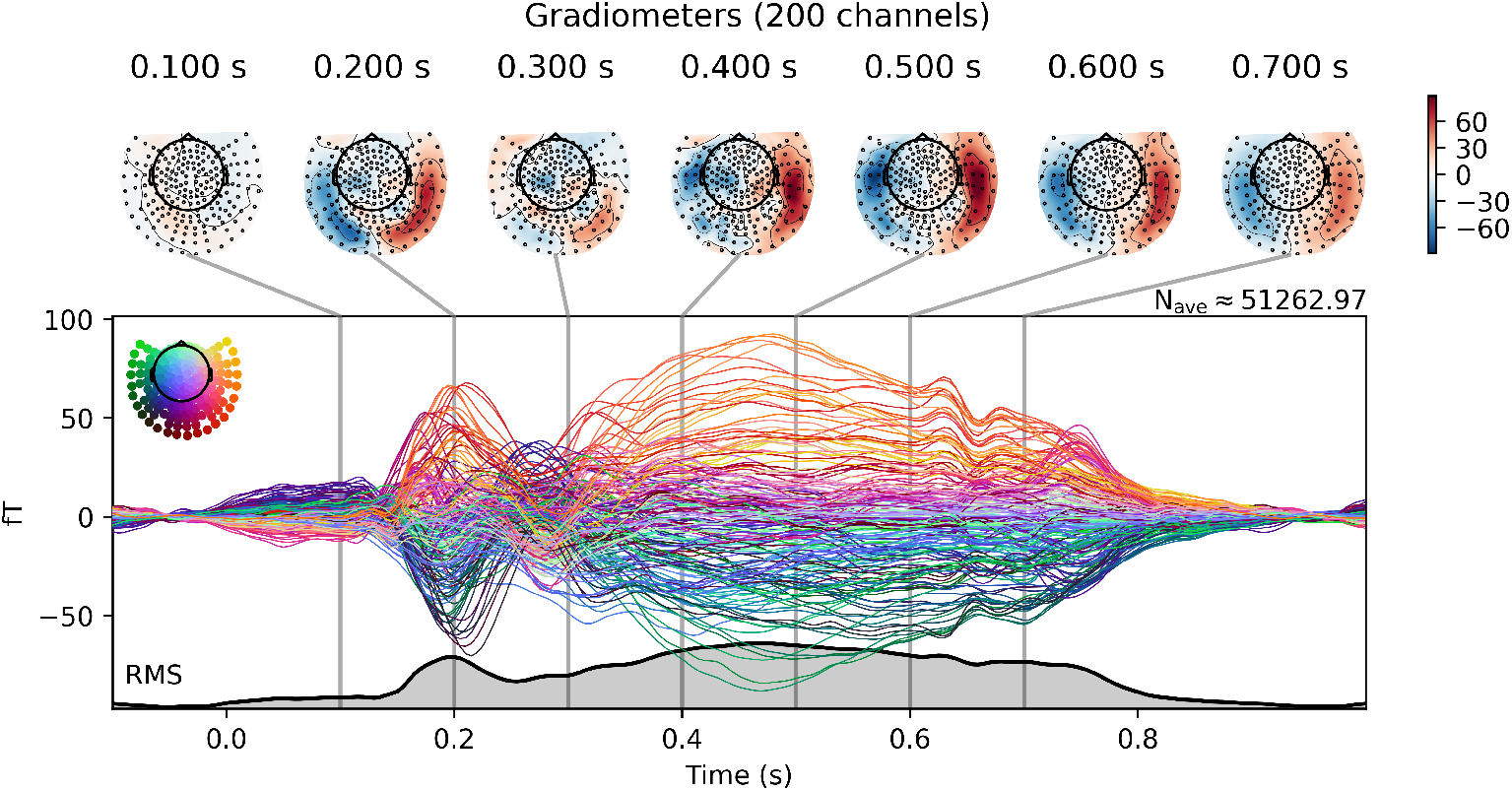
Average evoked response to all words across subjects (MEG).

#### EEG

As with the MEG data validation, the grand average is shown in Figure 5, where epochs spanned -0.1 s to 1.0 s with baseline correction applied over the prestimulus interval (-0.1 s to 0.0 s). See Preprocessing section for the details.

**Figure 5.**
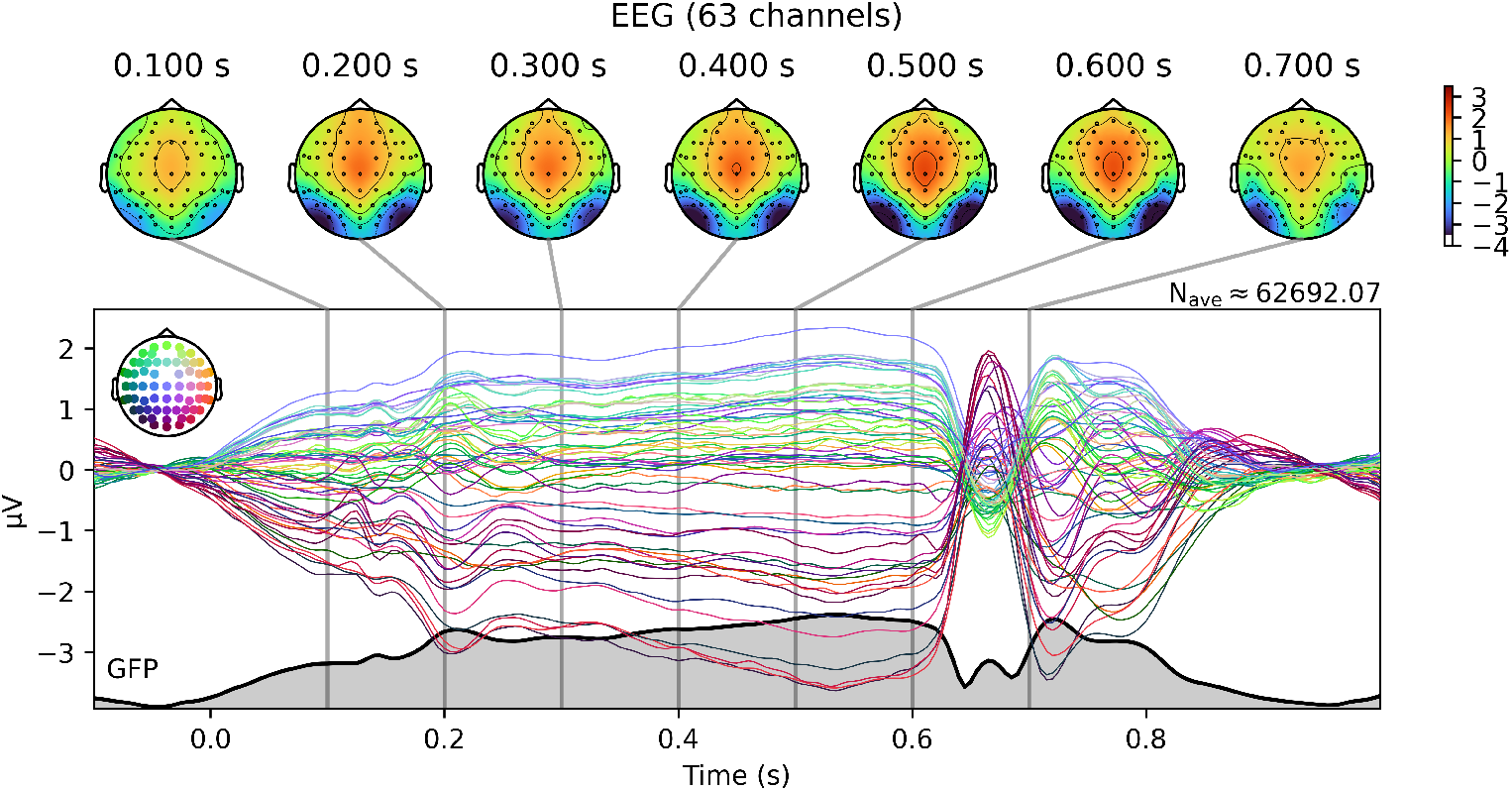
Average evoked response to all words across subjects (EEG).

### Usage Notes

Although we made the BCCWJ-Brain dataset publicly available, BCCWJ texts are not open source. To access the texts, please obtain a BCCWJ license; specifically, a contract for the Paid edition of the BCCWJ is required, as described at https://clrd.ninjal.ac.jp/bccwj/en/subscription.html. The script to reconstruct the text in the event files is provided on the BCCWJ platform (https://bccwj-data.ninjal.ac.jp/).

## Code Availability

All preprocessing scripts and validation analysis code are available at https://github.com/osekilab/BCCWJ-Brain.

## Acknowledgements

This work was supported by JSPS KAKENHI Grant Numbers JP19H05589, JP24H00085, JP24H00087, JST JPMJPR21C2, JST CREST JPMJCR2565, JST BOOST JPMJBY24B2; and the Yotta Informatics Project at Tohoku University, Japan.

## Author contributions statement

M.A., M.K., and Y.O. designed the experiments. Y.S., H.J., and A.K. collected the data for MEG and fMRI. Y.S. collected EEG data. Y.S. preprocssed and validated the data. Y.S. wrote the manuscript.

## Competing interests

No conflict of interests were reported by the authors.

## Footnotes

1 https://github.com/rordenlab/dcm2niix.

2 https://mne.tools/stable/index.html

3 https://eelbrain.readthedocs.io/en/stable/

4 Nilearn contributors, https://nilearn.github.io/stable/index.html

## Notes

### Competing Interest Statement

The authors have declared no competing interest.

## References

Kristijan Armeni, Umut Güçlü, Marcel van Gerven, and Jan-Mathijs Schoffelen. A 10-hour within-participant magnetoencephalography narrative dataset to test models of language comprehension. Scientific Data, 9(1):278, 2022. doi: 10.1038/s41597-022-01382-7.

Julia Berezutskaya, Mariska J. Vansteensel, Erik J. Aarnoutse, Zachary V. Freudenburg, Giovanni Piantoni, Mariana P. Branco, and Nick F. Ramsey. Open multimodal iEEG-fMRI dataset from naturalistic stimulation with a short audiovisual film. Scientific Data, 9(1):91, 2022. doi: 10.1038/s41597-022-01173-0. URL https://doi.org/10.1038/s41597-022-01173-0.

Shohini Bhattasali, Jonathan Brennan, Wen-Ming Luh, Berta Franzluebbers, and John Hale. The alice datasets: fMRI & EEG observations of natural language comprehension. In Proceedings of the Twelfth Language Resources and Evaluation Conference, pages 120–125, Marseille, France, May 2020. European Language Resources Association. ISBN 979-10-95546-34-4.

Jonathan R. Brennan, Yuval Nir, Uri Hasson, Rafael Malach, David J. Heeger, and Liina Pylkkänen. Syntactic structure building in the anterior temporal lobe during natural story listening. Brain and Language, 120(2):163–173, 2012. doi: 10.1016/j.bandl.2010.04.002.

Christian Brodbeck, Proloy Das, Marlies Gillis, Joshua P Kulasingham, Shohini Bhattasali, Phoebe Gaston, Philip Resnik, and Jonathan Z Simon. Eelbrain, a python toolkit for time-continuous analysis with temporal response functions. eLife, 12:e85012, nov 2023. ISSN 2050-084X. doi: 10.7554/eLife.85012. URL https://doi.org/10.7554/eLife.85012.

Charlotte Caucheteux, Alexandre Gramfort, and Jean-Rémi King. Evidence of a predictive coding hierarchy in the human brain listening to speech. Nature Human Behaviour, 7(3):430–441, 2023. doi: 10.1038/s41562-022-01516-2.

Cheng-Yeh Chen and Raghupathy Sivakumar. The mind’s transformer: Computational neuroanatomy of LLM-brain alignment. In The Fourteenth International Conference on Learning Representations, 2026. URL https://openreview.net/forum?id=PgIlCCNxdB.

Janice Chen, Yuan Chang Leong, Christopher J Honey, Chung H Yong, Kenneth A Norman, and Uri Hasson. Shared memories reveal shared structure in neural activity across individuals. Nature Neuroscience, 20(1):115–125, 2017. doi: 10.1038/nn.4450. URL https://doi.org/10.1038/nn.4450.

Nilearn contributors. nilearn. URL https://github.com/nilearn/nilearn.

Changjiang Gao, Zhengwu Ma, Jiajun Chen, Ping Li, Shujian Huang, and Jixing Li. Increasing alignment of large language models with language processing in the human brain. Nature Computational Science, 5(11):1080–1090, 2025. doi: 10.1038/s43588-025-00863-0. URL https://doi.org/10.1038/s43588-025-00863-0.

Ariel Goldstein, Zaid Zada, Eliav Buchnik, Mariano Schain, Amy Price, Bobbi Aubrey, Samuel A. Nastase, Amir Feder, Dotan Emanuel, Alon Cohen, Aren Jansen, Harshvardhan Gazula, Gina Choe, Aditi Rao, Catherine Kim, Colton Casto, Lora Fanda, Werner Doyle, Daniel Friedman, Patricia Dugan, Lucia Melloni, Roi Reichart, Sasha Devore, Adeen Flinker, Liat Hasenfratz, Omer Levy, Avinatan Hassidim, Michael Brenner, Yossi Matias, Kenneth A. Norman, Orrin Devinsky, and Uri Hasson. Shared computational principles for language processing in humans and deep language models. Nature Neuroscience, 25(3):369–380, 2022. doi: 10.1038/s41593-022-01026-4. URL https://doi.org/10.1038/s41593-022-01026-4.

Krzysztof J. Gorgolewski, Tibor Auer, Vince D. Calhoun, R. Cameron Craddock, Samir Das, Eugene P. Duff, Guillaume Flandin, Satrajit S. Ghosh, Tristan Glatard, Yaroslav O. Halchenko, Daniel A. Handwerker, Michael Hanke, David Keator, Xiangrui Li, Zachary Michael, Camille Maumet, B. Nolan Nichols, Thomas E. Nichols, John Pellman, Jean-Baptiste Poline, Ariel Rokem, Gunnar Schaefer, Vanessa Sochat, William Triplett, Jessica A. Turner, Gaël Varoquaux, and Russell A. Poldrack. The brain imaging data structure, a format for organizing and describing outputs of neuroimaging experiments. Scientific Data, 3(1):160044, 2016. doi: 10.1038/sdata.2016.44. URL https://doi.org/10.1038/sdata.2016.44.

Alexandre Gramfort, Martin Luessi, Eric Larson, Denis A. Engemann, Daniel Strohmeier, Christian Brodbeck, Roman Goj, Mainak Jas, Teon Brooks, Lauri Parkkonen, and Matti Hämäläinen. Meg and eeg data analysis with mne-python. Frontiers in Neuroscience, Volume 7-2013, 2013. ISSN 1662-453X. doi: 10.3389/fnins.2013.00267. URL https://www.frontiersin.org/journals/neuroscience/articles/10.3389/fnins.2013.00267.

Omer Faruk Gulban, Dylan Nielson, John Lee, Russ Poldrack, Chris Gorgolewski, Vanessasaurus, and Chris Markiewicz. poldracklab/pydeface: Pydeface v2.0.2, July 2022. URL 10.5281/zenodo.6856482.

Laura Gwilliams, Graham Flick, Alec Marantz, Liina Pylkkänen, David Poeppel, and Jean-Rémi King. Introducing meg-masc a high-quality magneto-encephalography dataset for evaluating natural speech processing. Scientific Data, 10(1):862, 2023. URL 10.1038/s41597-023-02752-5.

Sreejan Kumar, Theodore R. Sumers, Takateru Yamakoshi, Ariel Goldstein, Uri Hasson, Kenneth A. Norman, Thomas L. Griffiths, Robert D. Hawkins, and Samuel A. Nastase. Shared functional specialization in transformer-based language models and the human brain. Nature Communications, 15(1):5523, 2024. doi: 10.1038/s41467-024-49173-5.

Mathis Lamarre, Catherine Chen, and Fatma Deniz. Attention weights accurately predict language representations in the brain. In Yoav Goldberg, Zornitsa Kozareva, and Yue Zhang, editors, Findings of the Association for Computational Linguistics: EMNLP 2022, pages 4513–4529, Abu Dhabi, United Arab Emirates, December 2022. Association for Computational Linguistics. doi: 10.18653/v1/2022.findings-emnlp.330. URL https://aclanthology.org/2022.findings-emnlp.330/.

Amanda LeBel, Lauren Wagner, Shailee Jain, Aneesh Adhikari-Desai, Bhavin Gupta, Allyson Morgenthal, Jerry Tang, Lixiang Xu, and Alexander G. Huth. A natural language fmri dataset for voxelwise encoding models. Scientific Data, 10(1):555, 2023.

Jixing Li, Shohini Bhattasali, Shulin Zhang, Berta Franzluebbers, Wen-Ming Luh, R. Nathan Spreng, Jonathan R. Brennan, Yiming Yang, Christophe Pallier, and John T. Hale. Le petit prince multilingual naturalistic fmri corpus. Scientific Data, 9(1):530, 2022. doi: 10.1038/s41597-022-01625-7. URL https://doi.org/10.1038/s41597-022-01625-7.

Kikuo Maekawa, Makoto Yamazaki, Toshinobu Ogiso, Takehiko Maruyama, Hideki Ogura, Wakako Kashino, Hanae Koiso, Masaya Yamaguchi, Makiro Tanaka, and Yasuharu De. Balanced corpus of contemporary written japanese. Language Resrouces & Evaluation, 48:345–371, 2014.

Mohammad Momenian, Zhengwu Ma, Shuyi Wu, Chengcheng Wang, Jonathan Brennan, John Hale, Lars Meyer, and Jixing Li. Le petit prince hong kong (LPPHK): Naturalistic fMRI and EEG data from older cantonese speakers. Scientific Data, 11(1):992, 2024. doi: 10.1038/s41597-024-03745-8. URL https://doi.org/10.1038/s41597-024-03745-8.

Samuel A Nastase, Valeria Gazzola, Uri Hasson, and Christian Keysers. Measuring shared responses across subjects using intersubject correlation. Social Cognitive and Affective Neuroscience, 14(6): 667–685, 08 2019. ISSN 1749-5016. doi: 10.1093/scan/nsz037. URL https://doi.org/10.1093/scan/nsz037.

Samuel A. Nastase, Yun-Fei Liu, Hanna Hillman, Asieh Zadbood, Liat Hasenfratz, Neggin Keshavarzian, Janice Chen, Christopher J. Honey, Yaara Yeshurun, Mor Regev, Mai Nguyen, Claire H. C. Chang, Christopher Baldassano, Olga Lositsky, Erez Simony, Michael A. Chow, Yuan Chang Leong, Paula P. Brooks, Emily Micciche, Gina Choe, Ariel Goldstein, Tamara Vanderwal, Yaroslav O. Halchenko, Kenneth A. Norman, and Uri Hasson. The “narratives”fmri dataset for evaluating models of naturalistic language comprehension. Scientific Data, 8(1):250, 2021. doi: 10.1038/s41597-021-01033-3. URL https://doi.org/10.1038/s41597-021-01033-3.

Guiomar Niso, Krzysztof J. Gorgolewski, Elizabeth Bock, Teon L. Brooks, Guillaume Flandin, Alexandre Gramfort, Richard N. Henson, Mainak Jas, Vladimir Litvak, Jeremy T. Moreau, Robert Oostenveld, Jan-Mathijs Schoffelen, Francois Tadel, Joseph Wexler, and Sylvain Baillet. MEG-BIDS, the brain imaging data structure extended to magnetoencephalography. Scientific Data, 5(1): 180110, 2018. doi: 10.1038/sdata.2018.110. URL https://doi.org/10.1038/sdata.2018.110.

Jonathan W. Peirce. Psychopy—psychophysics software in python. Journal of Neuroscience Methods, 162(1):8–13, 2007. ISSN 0165-0270. doi: 10.1016/j.jneumeth.2006.11.017. URL https://www.sciencedirect.com/science/article/pii/S0165027006005772.

Jonathan W. Peirce. Generating stimuli for neuroscience using psychopy. Frontiers in Neuroinformatics, 2, 2009. ISSN 1662-5196. doi: 10.3389/neuro.11.010.2008. URL https://www.frontiersin.org/journals/neuroinformatics/articles/10.3389/neuro.11.010.2008.

Cyril R. Pernet, Stefan Appelhoff, Krzysztof J. Gorgolewski, Guillaume Flandin, Christophe Phillips, Arnaud Delorme, and Robert Oostenveld. EEG-BIDS, an extension to the brain imaging data structure for electroencephalography. Scientific Data, 6(1):103, 2019. doi: 10.1038/s41597-019-0104-8. URL https://doi.org/10.1038/s41597-019-0104-8.

Liina Pylkkänen and Alec Marantz. Tracking the time course of word recognition with meg. Trends in Cognitive Sciences, 7(5):187–189, 2003. ISSN 1364-6613. doi: 10.1016/S1364-6613(03)00092-5. URL https://www.sciencedirect.com/science/article/pii/S1364661303000925.

Jan-Mathijs Schoffelen, Robert Oostenveld, Nietzsche H. L. Lam, Julia Uddén, Annika Hultén, and Peter Hagoort. A 204-subject multimodal neuroimaging dataset to study language processing. Scientific Data, 6(1):17, 2019. doi: 10.1038/s41597-019-0020-y. URL https://doi.org/10.1038/s41597-019-0020-y.

Martin Schrimpf, Idan Asher Blank, Greta Tuckute, Carina Kauf, Eghbal A. Hosseini, Nancy Kanwisher, Joshua B. Tenenbaum, and Evelina Fedorenko. The neural architecture of language: Integrative modeling converges on predictive processing. Proceedings of the National Academy of Sciences, 118(45):e2105646118, 2021. doi: 10.1073/pnas.2105646118.

Qixuan Wang, Qian Zhou, Zhengwu Ma, Nan Wang, Tianyu Zhang, Yaoyao Fu, and Jixing Li. Le petit prince (lpp) multi-talker: Naturalistic 7T fMRI and EEG dataset. Scientific Data, 12(1):829, 2025. doi: 10.1038/s41597-025-05158-7. URL https://doi.org/10.1038/s41597-025-05158-7.

Shaonan Wang, Xiaohan Zhang, Jiajun Zhang, and Chengqing Zong. A synchronized multimodal neuroimaging dataset for studying brain language processing. Scientific Data, 9(1):590, 2022. doi: 10.1038/s41597-022-01708-5. URL https://doi.org/10.1038/s41597-022-01708-5.

